# Distinct and complementary mechanisms of oscillatory and aperiodic alpha activity in visuospatial attention

**DOI:** 10.1101/2025.03.20.644419

**Authors:** Runhao Lu, Elizabeth Pollitt, Alexandra Woolgar

## Abstract

Alpha oscillations are thought to play a key role in visuospatial attention, particularly through lateralisation mechanisms. However, whether this function is driven purely by oscillatory activity or also involves aperiodic neural components remains unclear, making it difficult to develop precise theoretical models of alpha function and attention. Using EEG and concurrent TMS-EEG, this study aimed to (1) disentangle the contributions of oscillatory and aperiodic alpha activity to visuospatial attention and (2) examine their causal roles by differentially modulating aperiodic and oscillatory components. First, across three independent EEG datasets, we found that both oscillatory and aperiodic responses in the alpha band contribute to spatial attention encoding and univariate lateralisation effects. The two signals were uncorrelated across electrodes and their combination yielded stronger effects than either signal separately, suggesting that they may play complementary roles. Then, we used concurrent TMS-EEG to modulate the two signals. Compared to arrhythmic TMS (ar-TMS), rhythmic TMS (rh-TMS), enhanced oscillatory alpha power, especially at the stimulated area, while decreasing aperiodic alpha power across the scalp. Despite these opposing effects, rh-TMS improved visuospatial attention representation carried by both oscillatory and aperiodic alpha signals, suggesting that both signals may support attentional processing through different mechanisms. Moreover, TMS-induced changes in oscillatory and aperiodic alpha decoding differentially predicted behavioural performance, with TMS-induced changes in oscillatory alpha decoding correlating with response errors and changes in aperiodic alpha decoding correlating with response speed. Together these findings reveal a functional dissociation between oscillatory and aperiodic activity in the alpha band. We suggest a dual mechanism for alpha band activity in supporting visuospatial attention, where the two components have distinct but complementary roles. Oscillatory components may primarily support attentional filtering and target prioritization, while aperiodic components may reflect overall neural excitability and cognitive efficiency. Both of these mechanisms contribute to successful visuospatial attention.

## Introduction

Visuospatial attention is a core cognitive function that allows us to prioritize information from relevant parts of the visual field while filtering out distracting information [1, 2]. This process has long been associated with posterior alpha oscillations (8–13 Hz), particularly through lateralisation mechanisms [3–9]. When attention is directed to one hemifield, posterior alpha power increases ipsilaterally and decreases contralaterally, a pattern thought to reflect functional inhibition of task-irrelevant brain regions and facilitation of relevant sensory processing [10–12].

However, changes in alpha power do not exclusively reflect oscillatory dynamics. Instead, they arise from a combination of oscillatory and aperiodic neural components, often referred to as a mixed signal (Figure 1A). Traditional analyses typically measure raw alpha power change, which can be easily confounded by aperiodic broadband power shifts or spectral rotations rather than reflecting genuine oscillatory modulation [13, 14]. This raises an important question: is the well-established role of alpha activity in encoding visuospatial attention purely oscillatory, or does aperiodic activity also contribute?

**Figure 1.**
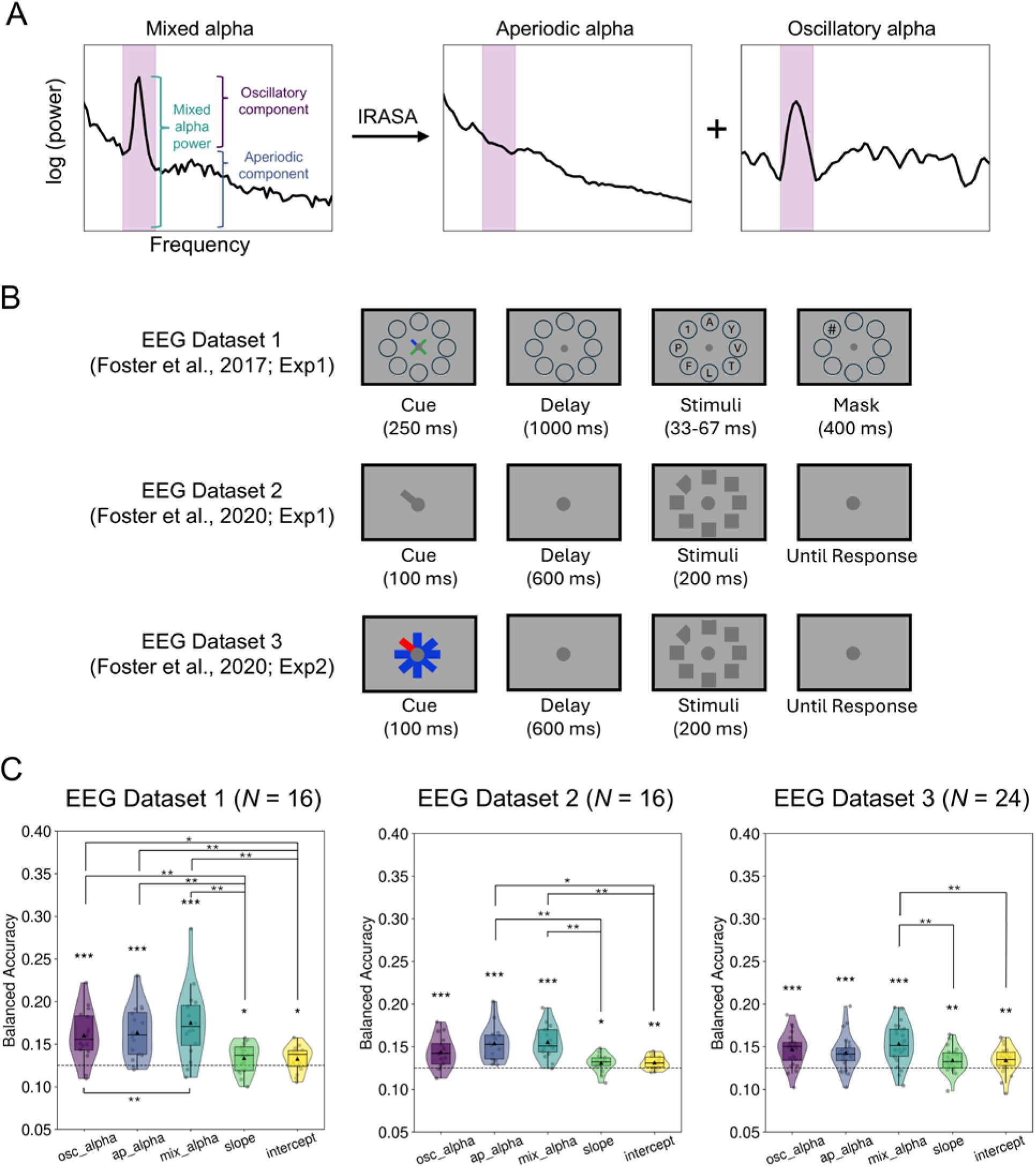
Oscillatory and aperiodic EEG activity encode visuospatial attention. (A) Illustration of mixed alpha activity, which encompasses both an oscillatory alpha peak and aperiodic broadband components. By applying methods such as IRASA, the mixed alpha signal can be decomposed into its separate aperiodic and oscillatory components. The purple shading highlights the 8-13 Hz alpha band. (B) Experimental designs of the three EEG datasets used in the study. In Dataset 1 (Foster et al., 2017, Exp 1; [33]), participants performed a covert visuospatial attention task, where they were cued to attend to one of eight locations. After a delay, they viewed a briefly presented target digit among distractors and had to identify the target digit using a number pad. In Datasets 2 and 3 (Foster et al., 2020, Exp 1 and Exp 2; [34]), participants were cued to a target location and, after a delay, had to identify whether the target (a diamond), presented among distractors (squares), was missing a left or right corner via keypress. Dataset 2 and 3 differed only in their cueing methods, with Dataset 3 using a visually balanced cue to control for asymmetries. (C) Balanced decoding accuracy for decoding visuospatial attention using five EEG signals from all EEG sensors for the three datasets (chance level: 0.125). Each dot represents one participant. The small triangle inside each violin plot indicates the mean, while the horizontal line represents the median. osc_alpha: oscillatory alpha power, ap_alpha: aperiodic alpha power, mix_alpha: mixed alpha power. * *p* _FDR_ < 0.05, ** *p* _FDR_ < 0.01, *** *p* _FDR_ < 0.001.

Unlike oscillatory activity, which reflects rhythmic fluctuations at specific frequencies, aperiodic activity represents the broadband characteristics of the neural signal, typically characterized by broadband power (including that within the alpha range), slope, and intercept. These aperiodic features are thought to reflect fundamental aspects of cortical dynamics such as neuronal excitability, population-level spiking activity, and/or excitation-inhibition (E/I) balance [15–17]. An increasing number of recent studies have emphasized the functional role of aperiodic activity in various cognitive processes, including attention and cognitive control [13, 14, 17–24]. For example, aperiodic slope is often suggested to flatten during the working memory maintenance period, potentially reflecting shifts in E/I balance that support working memory processes [13, 25]. Also, aperiodic broadband power has been shown to decrease with increasing cognitive control demands [13, 25, 26], suggesting a potential role in attentional processing. These findings indicate that aperiodic activity may not merely be background noise but a distinct and integral neural component supporting cognitive function, operating alongside oscillatory activity in a complementary manner.

While the role of oscillatory and aperiodic activity in cognition is increasingly recognized, further exploration requires causal manipulation to establish their distinct contributions. Non-invasive brain stimulation techniques, such as transcranial magnetic stimulation (TMS) and transcranial alternating current stimulation (tACS), provide powerful tools for modulating neural activity and assessing its influence on cognition. It has been well-established that rhythmic-TMS (rh-TMS) or rhythmic-tACS can entrain neural oscillations at specific frequencies, enhancing their power and inter-trial phase coherence [27–32]. However, whether aperiodic activity can be differentially modulated remains largely unexplored. Given its scale-free nature, aperiodic activity cannot be entrained in the same manner as oscillatory rhythms, suggesting that alternative stimulation approaches, such as arrhythmic TMS (ar-TMS), which delivers temporally irregular stimulation patterns, may be better suited for modulating aperiodic dynamics.

In this study, we first aimed to investigate whether the well-established role of alpha activity in visuospatial attention is purely driven by oscillatory dynamics or if aperiodic activity also plays a functional role. To address this, we analysed three open EEG datasets [33, 34], and applied irregular-resampling auto-spectral analysis (IRASA) [35] to separate the oscillatory and aperiodic EEG components. Then, we used multivariate decoding and lateralisation measures to assess their respective contributions to supporting visuospatial attention. Second, we aimed to assess the causal influence of oscillatory and aperiodic activity on visuospatial attention. To achieve this, we re-analysed our recent TMS-EEG dataset [32] in which participants performed a selective attention task while receiving either rh-TMS at their individual alpha frequency (IAF) or ar-TMS. This allowed us to investigate whether rh-TMS selectively enhanced oscillatory alpha power while differentially modulating aperiodic activity, relative to ar-TMS, and how these neural modulations influence both the representation of visuospatial attention and associated behavioural performance.

## Results

### Both oscillatory and aperiodic activity encode visuospatial attention

The first objective of this study was to examine whether visuospatial attention is encoded in purely oscillatory activity, aperiodic activity, or both. To address this, we applied multivariate decoding analysis to three independent EEG datasets that involved covert visuospatial attention tasks (Figure 1B; Dataset 1: Experiment 1 from Foster, et al. [33]; Dataset 2 and Dataset 3: Experiment 1 and 2 from Foster, et al. [34]). To isolate oscillatory and aperiodic contributions, we first removed event-related potentials (i.e., the average time-locked signal across trials) from each trial for each condition and then applied the IRASA method [35] to decompose EEG signals into their oscillatory and aperiodic components. From these we extracted five key signal types for decoding analysis: Oscillatory alpha activity (8-13 Hz), aperiodic alpha activity (8-13 Hz of the aperiodic broadband power), mixed alpha activity (i.e., combination of both oscillatory and aperiodic activity at 8-13 Hz), aperiodic slope and aperiodic intercept. For all three datasets, we performed decoding analysis using signals from all EEG sensors during the delay period. We trained a support vector machine (SVM) classifier to decode the cued spatial location in an 8-way classification from each of these signal types.

As illustrated in Figure 1C, across all three datasets, we found that all five signals of interest could decode spatial attention information. Specifically, oscillatory alpha, aperiodic alpha, and mixed alpha activity exhibited strong decoding performance (4.05< *t*s < 6.36, *p*s _FDR_ < 0.001), while aperiodic slope and intercept showed significant but relatively weaker decoding performance (1.97 < *t*s < 3.29; *p*s _FDR_ < 0.04). Numerically, mixed alpha activity had the highest decoding performance, suggesting that combining oscillatory and aperiodic contributions may enhance the performance to encode visuospatial attention by providing complimentary information.

To formally compare decoding performance between different signals of interest, we performed a one-way repeated-measures ANOVA for each dataset. Results showed a significant main effect of signal type in all three datasets [all *F*s > 6.32; *p*s < 0.001]. Post-hoc paired comparisons (Figure 1C) indicated that aperiodic slope and intercept showed significantly lower decoding accuracy compared to one or more alpha-band signals (i.e., oscillatory, aperiodic, or mixed alpha).

Thus, across three independent EEG datasets we found that the location of visuospatial attention (where a person is attending) can be decoded consistently from both oscillatory and aperiodic activity in the alpha band. Among the signal types examined, mixed alpha activity showed the strongest decoding performance numerically, potentially due to its integration of oscillatory and aperiodic contributions. Oscillatory and aperiodic alpha activity ranked next, both demonstrating strong decoding performance. In contrast, while aperiodic slope and intercept also contained spatial information, their decoding accuracy was significantly lower than that of alpha-band signals.

### Lateralised alpha modulation arises from both oscillatory and aperiodic activity

Given that all five signals encoded visuospatial attention, we next investigated whether they contributed to the classic lateralisation effect observed during spatial attention. First, we examined the topographic distributions of lateralisation effects by subtracting the attend-right condition (3 o’clock) from the attend-left condition (9 o’clock). As shown in Figure 2A, across all three EEG datasets, mixed alpha activity again appeared to exhibit the strongest lateralisation effects, followed by oscillatory alpha activity. Aperiodic activities demonstrated relatively weak lateralisation overall, though aperiodic alpha and slope showed some degree of lateralisation in Dataset 2.

**Figure 2.**
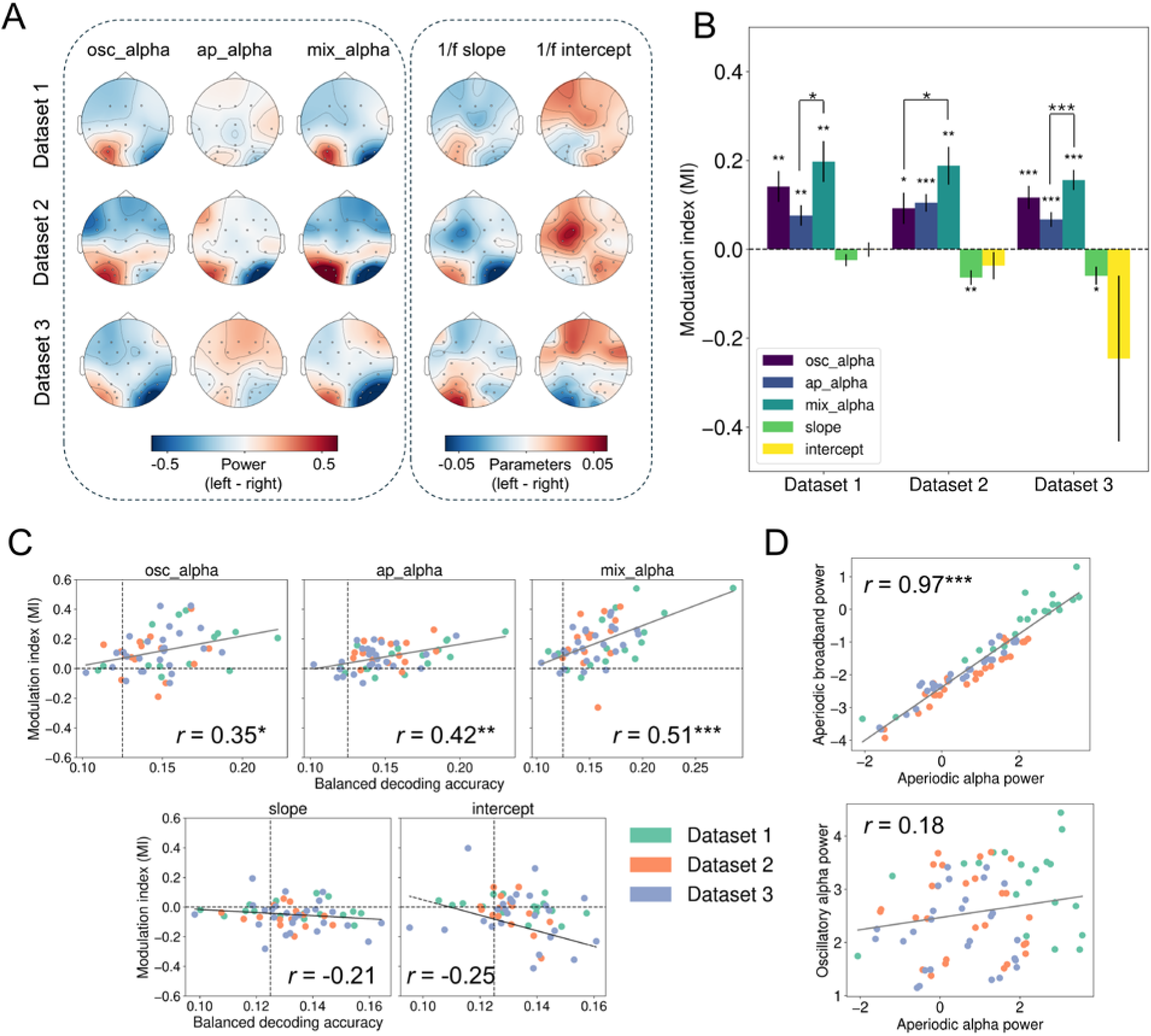
Lateralisation of alpha activity arises from both oscillatory and aperiodic components. (A) Topographic maps of lateralisation effects for each EEG signal across the three datasets. Lateralisation was computed by subtracting the attend-right condition (3 o’clock) from the attend-left condition (9 o’clock). (B) Modulation index (MI) quantifies lateralisation effects across datasets. Significance markings show differences from 0 and differences between oscillatory alpha, aperiodic alpha, and mixed alpha (we did not make statistical comparisons with slope and intercept). Error bars show standard errors of the mean. (C) Spearman’s correlations between MI and decoding accuracy across participants. Data were pooled across the three datasets, with different colours indicating different datasets. For the intercept plot, one outlier (MI = -4.92, decoding = 0.14) was not included for visual clarity. (D) Spearman’s correlations between aperiodic alpha power and aperiodic broadband power (3-30 Hz; top plot) and oscillatory alpha power (bottom) across electrodes. Data were pooled across three datasets, with different colours indicating different datasets. * *p* _FDR_ < 0.05, ** *p* _FDR_ < 0.01, *** *p* _FDR_ < 0.001.

To quantitatively assess the lateralisation effects, we computed the modulation index (MI; see Methods) for each signal within each dataset (Figure 2B). Results showed that mixed alpha activity had the highest MI across all three datasets, followed by oscillatory alpha activity and aperiodic alpha activity. This suggested that oscillatory and aperiodic alpha activity may make complementary contributions to lateralisation effects. Aperiodic slope and intercept displayed inconsistent lateralisation patterns across datasets—while slope showed significant lateralisation in Dataset 2 and 3, intercept did not exhibit any significant lateralisation effects in any dataset. Notably, MI values are typically expected to be positive, as spectral power is inherently positive. However, the negative MI values for aperiodic slope and intercept may stem from their inherent sign differences (e.g., aperiodic slope is always negative). In Datasets 2 and 3, when participants attended to the left, their aperiodic slope flattened in the ipsilateral hemisphere while steepened in the contralateral hemisphere.

To statistically compare lateralisation across different signals, we performed a one-way repeated-measures ANOVA for each dataset, focusing on oscillatory alpha, aperiodic alpha, and mixed alpha (excluding aperiodic slope and intercept due to their distinct scales and inconsistent lateralization patterns). ANOVA results confirmed that mixed alpha activity showed the strongest MI, significantly exceeding either oscillatory or aperiodic alpha activity in all three datasets (see Figure 2B).

Next, we asked whether lateralization effects were correlated with decoding performance. If lateralisation reflects a key neural signature of visuospatial attention, we would expect stronger lateralization to be associated with higher decoding accuracy. To test this, we pooled data across all three datasets (*N* = 56) and performed Spearman’s correlations between MI and decoding performance across participants for each signal (Figure 2C). We found significant positive correlations between MI and decoding performance for oscillatory alpha (*r* = 0.35, *p* _FDR_ = 0.01), aperiodic alpha (*r* = 0.42, *p* _FDR_ = 0.003), and mixed alpha activity (*r* = 0.51, *p* _FDR_ < 0.001). However, no significant correlations were found for aperiodic slope or intercept (|*r|*s < 0.25, *p*s _FDR_ > 0.05) where MI and decoding accuracies were low.

In summary, across three independent EEG datasets, mixed alpha activity demonstrated the strongest lateralisation effects, and both oscillatory and aperiodic activities in alpha band also exhibited significant lateralisation. In contrast, aperiodic slope and intercept showed weak and inconsistent lateralisation across datasets. Moreover, we found significant positive correlations between lateralisation effects and decoding performance in oscillatory, aperiodic, and mixed alpha activity (but not in aperiodic slope and intercept), further underscoring their roles in encoding visuospatial attention.

### Aperiodic alpha activity reflects broadband power rather than oscillatory contamination

Since both aperiodic and oscillatory alpha activity contribute to the encoding of visuospatial attention and exhibit lateralisation effects, we carried out an additional analysis to establish the extent to which the two signals were distinct. To investigate this, we first averaged EEG activity across participants for each signal and each dataset. We then concatenated the data across all three datasets and computed Spearman’s correlations between aperiodic alpha power and both aperiodic broadband power (3-30 Hz) and oscillatory alpha power across electrodes (Figure 2D). There was an extremely strong correlation between aperiodic alpha and aperiodic broadband power (*r* = 0.97, *p* < 0.001), whereas its correlation with oscillatory alpha was much weaker and non-significant (*r* = 0.18, *p* = 0.11). These findings confirm that aperiodic alpha activity primarily reflects broadband aperiodic power rather than being influenced by oscillatory activity, reinforcing its distinct neural basis.

### Alpha rh-TMS and ar-TMS causally and differentially modulate oscillatory alpha and aperiodic broadband activity

Based on three independent EEG datasets, we have demonstrated that both oscillatory alpha and aperiodic alpha activity (reflecting aperiodic broadband power) play important roles in encoding visuospatial attention, likely through lateralisation mechanisms. Next, we aimed to establish causal evidence for their functional contributions, by differentially manipulating oscillatory and aperiodic activity using TMS. We analysed data from our recent concurrent TMS-EEG experiment [32] in which participants performed a visuospatial attention task under rhythmic-, arrhythmic-, and no-TMS conditions (Figure 3A). The rh-TMS condition involved five rhythmic TMS pulses at the IAF, delivered during the delay period after the spatial cue but before stimulus onset. In contrast, the ar-TMS condition delivered five pulses with an arrhythmic temporal structure, maintaining the same total duration as rh-TMS but disrupting its rhythmic structure by jittering the timing of pulses 2 3 and 4. Additionally, participants completed a no-TMS condition, in which no stimulation was applied during the delay period. As before, we separated EEG signals into the 5 signal types.

**Figure 3.**
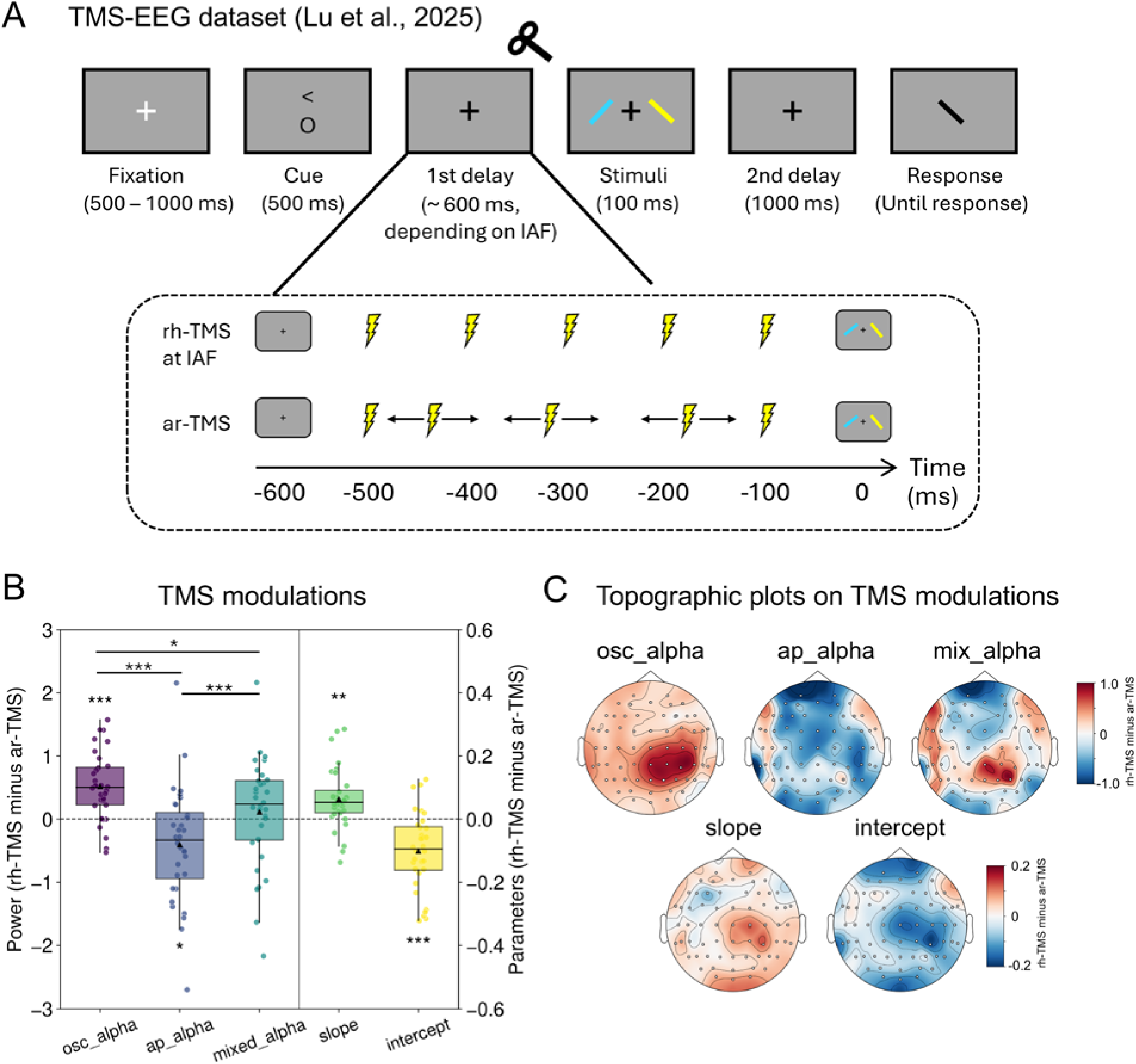
Rhythmic- and arrhythmic-TMS differentially modulate oscillatory and aperiodic neural activity. (A) Experimental design for the TMS-EEG dataset (Lu et al., 2025; [32]). Participants performed a selective attention task while receiving either rhythmic TMS (rh-TMS) or arrhythmic TMS (ar-TMS) at their individual alpha frequency (IAF). TMS was applied during the delay period before stimulus onset. No stimulation was delivered in the no-TMS condition. (B) TMS modulation effects of rh-TMS compared to ar-TMS during the delay period over right posterior regions. Box plots show activity differences (rh-TMS minus ar-TMS) for each type of signal. * *p* _FDR_ < 0.05, ** *p* _FDR_ < 0.01, *** *p* _FDR_ < 0.001 (C) Topographic distribution of TMS effects (rh-TMS minus ar-TMS) for each type of signal during the delay period.

We first examined the TMS modulation effects for each type of EEG signal, by subtracting activity in the ar-TMS condition from the rh-TMS condition during the delay period (-600 to 0 ms) around the stimulation location (right posterior areas; see Methods). As shown in Figure 3B, compared to ar-TMS, rh-TMS significantly increased oscillatory alpha power (*t* = 5.44, *p* _FDR_ < 0.001), while significantly decreasing aperiodic alpha (broadband) power (*t* = -2.53, *p* _FDR_ = 0.02). Mixed alpha power showed an intermediate response, numerically positive but not statistically different from zero (*t* = 0.70, *p* _FDR_ = 0.49). This seems likely to be because mixed alpha reflects both oscillatory and aperiodic components that showed opposing patterns and partially cancelled each other out. A repeated-measures ANOVA among the three signals confirmed a significant effect of signal type [*F* (2,62) = 15.83, *p* < 0.001]. Post-hoc comparisons showed that rh-TMS enhanced oscillatory alpha significantly more than aperiodic (*p* < 0.001) and mixed alpha (*p* = 0.02).

Topographic plots (Figure 3C) confirmed that rh-TMS-induced oscillatory alpha enhancement was localised to the stimulation area around the right parietal regions. In contrast, the relative suppression of aperiodic alpha (broadband) power induced by rh-TMS, compared to ar-TMS, was more widespread across the brain and was not restricted to the stimulation site. The mixed alpha modulation pattern replicated findings reported in [32], showing a mixture of TMS modulation for both oscillatory and aperiodic activity.

We also found that rh-TMS significantly modulated aperiodic parameters, specifically flattening the aperiodic slope (*t* = 3.67, *p* _FDR_ = 0.002) and decreasing the aperiodic intercept (*t* = -4.65, *p* _FDR_ < 0.001) (Figure 3B). The spatial distribution of these TMS-induced changes showed a combination of localised and more widespread effects, with some degree of specificity around the stimulation site but also modulation extending beyond it (Figure 3C). The direction and spatial distribution of aperiodic intercept modulation closely resembled that of aperiodic alpha (broadband) power, likely due to their inherent relationship.

We additionally compared the effect of rh-TMS and ar-TMS on each signal type relative to the no-TMS condition (Figure S1). As expected, rh-TMS (*t* = 3.17, *p* _FDR_ = 0.006) significantly increased oscillatory alpha power relative to no-TMS, whereas ar-TMS did not induce an increase (*t* = -1.40, *p* _FDR_ = 0.22). However, both rh-TMS and ar-TMS increased aperiodic alpha (*t* = 7.13 and 7.18, respectively, *p*s _FDR_ < 0.001) and mixed alpha power (*t* = 8.36 and 6.29, respectively, *p*s _FDR_ < 0.001) compared to no-TMS, with ar-TMS producing a significantly stronger effect than rh-TMS (Figure 3B). These results clarify that, while only rh-TMS selectively enhances oscillatory alpha, both TMS protocols modulate aperiodic broadband power, though to significantly different degrees.

These findings provide novel evidence that TMS can differentially modulate oscillatory and aperiodic activity. Compared to ar-TMS, rh-TMS specifically entrained oscillatory alpha power, especially at the stimulation site. In contrast, ar-TMS, relative to rh-TMS, increased aperiodic broadband power and intercept in a widespread manner. Aperiodic slope was also modulated by TMS, with rh-TMS relatively flattening the slope (i.e., ar-TMS relatively steepening it). These results suggest that it is promising to use rh-TMS and ar-TMS to differentially modulate oscillatory and aperiodic activity, providing opportunities to investigate their distinct causal roles in cognition, including visuospatial attention.

### Alpha rh-TMS improves neural representation of visuospatial attention through both oscillatory and aperiodic activity

Given that rh-TMS and ar-TMS differentially modulate oscillatory and aperiodic activity, we next investigated how these modulations influence the coding of visuospatial attention. We first applied multivariate decoding analysis to assess whether the five signals of interest encode visuospatial attention during the delay period under different TMS conditions (rh-TME and ar-TMS). As shown in Figure 4A, results replicated findings from the three EEG datasets (Figure 1C), demonstrating that all five signals, under both TMS conditions, significantly decoded visuospatial information with above-chance accuracy (all *p*s _FDR_ < 0.05).

**Figure 4.**
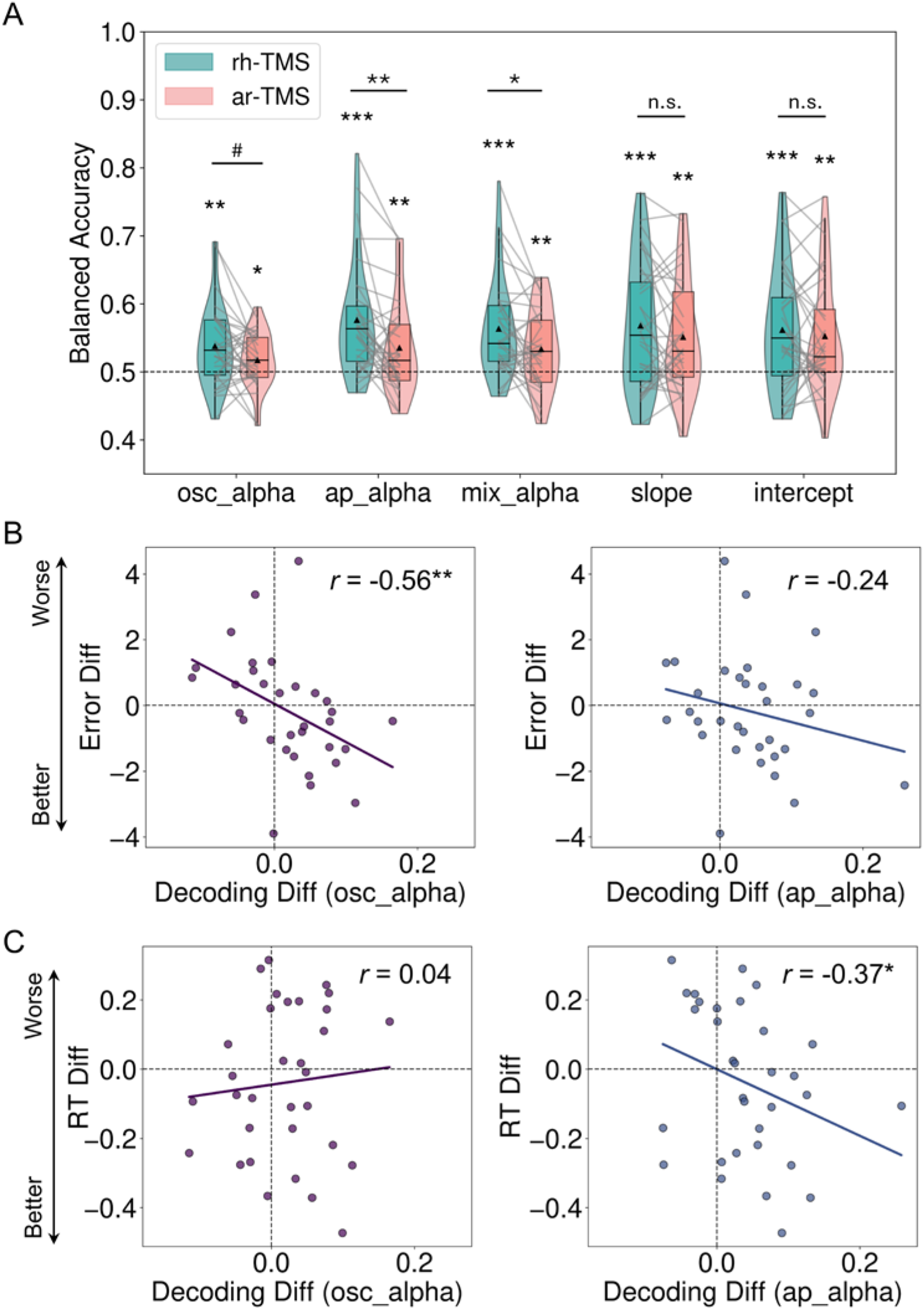
Rhythmic-TMS enhances neural representation of visuospatial attention through both oscillatory and aperiodic activity. (A) Balanced decoding accuracy for decoding visuospatial attention under rhythmic TMS (rh-TMS) and arrhythmic TMS (ar-TMS) conditions for each signal type (chance level: 0.5). Individual participant data are shown as grey lines connecting rh-TMS and ar-TMS conditions. (B) Correlations between TMS-induced changes in oscillatory and aperiodic alpha decoding and behavioural errors. Negative values on the y-axis (Error Diff) indicate fewer errors, meaning better performance, in the rh-TMS condition. (C) Correlations between TMS-induced changes in oscillatory and aperiodic alpha decoding and reaction times (RT). Negative values on the y-axis (RT Diff) indicate faster reaction times, meaning better performance, in the rh-TMS condition. ^#^ *p* _FDR_ < 0.10, * *p* _FDR_ < 0.05, ** *p* _FDR_ < 0.01, *** *p* _FDR_ < 0.001

To determine whether different TMS conditions influenced decoding performance, we conducted a 5 (signal type) × 2 (TMS condition) repeated-measures ANOVA. Results revealed a significant main effect of TMS condition [*F* (1,30) = 7.15, *p* = 0.01], with overall decoding performance being significantly higher in the rh-TMS condition compared to the ar-TMS condition. There was no significant interaction effect between signal type and TMS condition [*F* (4,120) = 1.79, *p* = 0.14], reflecting a consistent trend of TMS effect across the different signals. Exploratory paired t-tests showed that decoding accuracy was significantly higher under rh-TMS compared to ar-TMS for aperiodic alpha (broadband) power (*t* = 3.38, *p* _FDR_ = 0.005) and mixed alpha activity (*t* = 2.34, *p* _FDR_ = 0.03), and marginally higher under rh-TMS compared to ar-TMS for oscillatory alpha power (*t* = 1.72, *p* _FDR_ = 0.08). The differences were not significant for aperiodic slope or intercept-based decoding (*t*s < 1.23, *p*s _FDR_ > 0.14).

These findings suggest that, although rh-TMS enhances oscillatory alpha power and induces a smaller increase in aperiodic broadband power, compared to ar-TMS, it improves the neural representation of visuospatial attention through both oscillatory and aperiodic signals. The direction of this effect is particularly interesting for the aperiodic alpha component: while the overall level of aperiodic power is increased less by rh-TMS (Figure 3B), its multivariate signal (amplitude and distribution of aperiodic power over the scalp) also becomes more discriminatory between attend left and attend right conditions.

### TMS-induced decoding changes on oscillatory and aperiodic alpha activity differentially predict behavioural performance

Finally, to investigate the behavioural relevance of these differential TMS-induced neural modulations, we examined the relationship between changes in decoding accuracy and changes in behavioural performance (rh-TMS minus ar-TMS) using Spearman’s correlation. For oscillatory and aperiodic alpha activity separately, we correlated the TMS-induced differences in neural decoding with differences in behavioural errors and reaction times (RT).

As shown in Figure 4B, TMS-induced changes in oscillatory alpha decoding significantly and negatively correlated with changes in behavioural errors (*r* = -0.56, *p* _FDR_ = 0.002). This suggested that rh-TMS enhanced oscillatory coding of visuospatial attention was associated with improved behavioural precision. The correlation between TMS-induced aperiodic alpha decoding and behavioural errors was in the same direction, but not significant (*r* = -0.24, *p* _FDR_ = 0.13).

In contrast, as show in Figure 4C, TMS-induced decoding changes in *aperiodic* alpha activity was significantly and negatively correlated with RT changes (*r* = -0.37, *p* _FDR_ = 0.04). This indicated that rh-TMS enhanced coding of visuospatial attention in aperiodic alpha activity was linked to faster responses. However, there was no significant relationship between TMS-induced changes in oscillatory alpha decoding and RT (*r* = 0.04, *p* _FDR_ = 0.41).

These findings reveal a functional dissociation between oscillatory and aperiodic alpha (broadband) activity in behaviour. Oscillatory alpha activity predicts behavioural accuracy, likely reflecting a role in prioritising the process of the target while inhibiting distractors, whereas aperiodic alpha (broadband) activity is more closely tied to response speed, potentially reflecting global neural excitability and cognitive efficiency. Together, these results further highlight the distinct but complementary roles of oscillatory and aperiodic activity in visuospatial attention.

## Discussion

In this study, we investigated the distinct and complementary roles of oscillatory and aperiodic alpha activity in visuospatial attention using a combination of EEG and TMS-EEG experiments. Across three independent EEG datasets, we found that both oscillatory alpha and aperiodic alpha activity (reflecting aperiodic broadband power) encoded visuospatial attention, with mixed alpha activity showing the strongest decoding performance. Similarly, lateralisation effects, a hallmark of spatial attention, were present in both oscillatory and aperiodic alpha activity, again with mixed alpha showing the strongest lateralisation. In contrast, aperiodic slope and intercept exhibited weaker and more inconsistent decoding performance and lateralisation effects, indicating a more limited role in spatial attention. Using TMS-EEG, we further demonstrated that, compared to ar-TMS, rh-TMS at the IAF selectively enhanced oscillatory alpha power while inducing significantly less aperiodic broadband power. Despite these opposing effects, rh-TMS improved the neural representation of visuospatial attention through both oscillatory alpha and aperiodic broadband power. Finally, TMS-induced decoding changes in oscillatory and aperiodic alpha activity differentially predicted behavioural error and response speed, respectively. These findings provide causal evidence for the distinct yet collaborative contributions of oscillatory alpha and aperiodic activity to attentional processing.

It has been well established that alpha oscillations play a critical role in visuospatial attention through a lateralisation mechanism [3, 7, 9], facilitating processing in the attended hemifield while suppressing irrelevant information [10, 11]. However, most previous studies examining this effect have focused on mixed alpha activity, making it unclear which component drives this effect. Recent studies have emphasised that aperiodic components can significantly impact the interpretation of electrophysiological results and, if not properly accounted for, may lead to misconceptions in the theoretical frameworks of neural oscillations [13, 14, 17]. For instance, a recent study suggested that a substantial proportion of modulations previously attributed to frontal theta oscillations may instead be explained by changes in aperiodic activity [36]. These findings highlight the pressing need to revisit classic theories, including the role of alpha oscillations and lateralisation in visuospatial attention, to achieve a more precise understanding of alpha function and the neural mechanisms underlying selective attention.

By disentangling oscillatory and aperiodic alpha activity, we demonstrated that both of them contribute to visuospatial attention encoding and lateralisation. These results support the well-established role of oscillatory alpha activity in spatial attention, reinforcing previous findings that alpha power and lateralisation are functionally significant [10, 37]. At the same time, we show that aperiodic alpha activity also plays a role, suggesting that some prior effects attributed solely to oscillations may have involved aperiodic contributions. Notably, while oscillatory and aperiodic alpha power independently carried spatial attention information, their combined activity exhibited numerically stronger decoding accuracy and lateralisation effects, indicating that oscillatory and aperiodic alpha activity may complementarily support spatial attention encoding.

Importantly, we found that aperiodic alpha activity strongly reflects aperiodic broadband power, with an extremely high correlation (*r* = 0.97), but does not exhibit a significant relationship with oscillatory alpha power (*r* = 0.18). This finding confirms that oscillatory and aperiodic alpha activity are inherently distinct, yet both contribute to the neural representation of visuospatial attention.

Another key objective of this study was to causally and differentially manipulate oscillatory and aperiodic activity and to examine their distinct roles in visuospatial attention. It is well established that rh-TMS at a specific frequency can entrain neural oscillations in that band, leading to increased power and inter-trial phase coherence [27, 28, 32, 38, 39]. However, no studies have reported the successful manipulation of aperiodic activity using non-invasive brain stimulation like TMS. Given the aperiodic nature of ar-TMS, we tested whether ar-TMS could selectively enhance aperiodic activity relative to rh-TMS. Our findings revealed that rh-TMS specifically increased oscillatory alpha power compared to both ar-TMS and no-TMS conditions, especially at the stimulation site, reinforcing previous entrainment studies showing that rh-TMS effectively boosts pure oscillatory alpha activity. Notably, entrainment effects were stronger and clearer when analysing pure oscillatory power rather than mixed alpha power, highlighting the importance of isolating oscillatory components in neuromodulation studies. The reason for this was that although both rh-TMS and ar-TMS increased aperiodic broadband power relative to no-TMS condition, ar-TMS produced a significantly stronger increase in aperiodic broadband power than rh-TMS. Thus we found that oscillatory and aperiodic alpha activity can be differentially modulated using rh-TMS and ar-TMS, opening new possibilities for future research to causally manipulate aperiodic activity and investigate its functional role in cognition.

Aligning with our previous findings from time-resolved decoding [32], rh-TMS entrained oscillatory alpha activity and enhanced alpha-based decoding of spatial attention, reinforcing the functional role of anticipatory alpha oscillations in attentional selection. Interestingly, although rh-TMS *decreased* aperiodic alpha power compared to ar-TMS, it still *enhanced* the neural representation of spatial attention through aperiodic activity, compared to the same ar-TMS baseline. This finding further supports the notion that oscillatory and aperiodic alpha components may play distinct yet complementary roles in visuospatial attention. Recent research investigating the contributions of oscillatory and aperiodic activity to domain-general cognition [26] has shown that wide-distributed aperiodic broadband power decreases with increasing cognitive control demands. Perhaps, then, the lower level of aperiodic broadband power induced by rh-TMS in our study may be beneficial for cognitive control processes, including visuospatial attention.

Rather than serving as a gating or inhibitory control mechanism typically attributed to oscillatory alpha [10], aperiodic broadband power has been proposed to reflect fundamental neural mechanisms such as population-level spiking activity and the E/I balance [13, 17, 21, 40]. Thus, it is possible that oscillatory and aperiodic activity provide distinct but complementary mechanisms in supporting visuospatial attention. This functional dissociation was further supported by our correlation analysis, which revealed that TMS-induced changes in oscillatory and aperiodic alpha decoding were differentially linked to behavioural performance: oscillatory alpha decoding changes correlated with behavioural errors, whereas aperiodic alpha decoding changes were associated with response speed. These findings align with the proposal that oscillatory alpha may play a direct role in attentional filtering and inhibitory control, allowing for better suppression of distractors and more precise target selection [10]. In contrast, aperiodic activity may be more closely related to state-dependent neural excitability and cognitive efficiency, given its association with neuronal gain control, large-scale network dynamics, and E/I balance [15, 17]. The observed decrease in aperiodic power, alongside improved decoding and faster response times, suggests that lower aperiodic activity may reflect a more functionally distinct neural representation of attentional states, potentially allowing for more efficient processing. We suggest a dual mechanism of alpha-band activity in visuospatial attention, where oscillatory alpha may provide structured, periodic modulation to prioritize target processing, while aperiodic alpha activity (reflecting broadband aperiodic responses) may contribute to sustained network excitability and adaptive cognitive control.

In conclusion, this study provides novel evidence that oscillatory and aperiodic alpha activity play distinct but complementary roles in visuospatial attention. Using EEG and TMS-EEG, we demonstrated that both oscillatory and aperiodic alpha encode visuospatial attention and contribute to lateralisation effects, with mixed alpha activity showing the strongest effects. By employing rhythmic and arrhythmic TMS, we established a novel method for causally and differentially modulating oscillatory and aperiodic activity. While rh-TMS selectively enhanced oscillatory alpha power and induced significantly less aperiodic broadband activity relative to ar-TMS, it improved spatial attention representation through both neural components, highlighting their distinct yet collaborative contributions to attentional processing. Furthermore, TMS-induced changes in oscillatory and aperiodic alpha decoding were specifically linked to behavioural error and response speed, respectively, suggesting a potential functional dissociation in which oscillatory alpha may support attentional filtering whereas the amplitude and distribution of aperiodic activity may modulate broader attentional states and processing efficiency. These findings refine theories on the neural mechanisms of visuospatial attention and clarify the distinct but complementary roles of oscillatory and aperiodic alpha activity in visuospatial attention.

## Methods

### Participants

Across the three open EEG datasets [33, 34] and the TMS-EEG dataset [32], participants were between 18 and 36 years old, reported normal or corrected-to-normal visual acuity, and provided informed consent in accordance with the ethical guidelines of their respective institutions (University of Oregon for Dataset 1; University of Chicago for Datasets 2 and 3; University of Cambridge for the TMS-EEG dataset).

After excluding participants due to excessive artifacts or poor performance, the final samples included 16 participants (6 female, 10 male) for EEG Dataset 1 (Foster et al., 2017, Experiment 1; [33]), 16 participants (7 female, 9 male) for EEG Dataset 2 (Foster et al., 2020, Experiment 1; [34]), 26 participants (15 female, 11 male) for EEG Dataset 3 (Foster et al., 2020, Experiment 2; [34]), and 32 participants (18 female, 14 male) for the TMS-EEG dataset [32]. For analyses involving the no-TMS condition (Figure S1), a subset of 30 participants (18 female, 12 male) from the TMS-EEG dataset was included due to additional exclusions based on excessive artifacts. Full details on participant selection for these four datasets are reported in [33], [34], and [32], respectively.

### Experimental procedure

Across the three EEG datasets (Figure 1B; [33, 34]), participants performed visuospatial attention tasks that required attending to a cued location to identify a target among distractors. For EEG Dataset 1 ([33], Experiment 1), each trial began with a central cue (a cross with one coloured arm) for 250 ms, directing attention to one of eight locations (87.5% valid). After a delay period (1000 ms), a target digit appeared among distractor letters for a brief duration (33-67 ms) before being masked with a pound sign. Participants were required to identify the target digit. We analysed both valid- and invalid-cue trials as both conditions engaged visuospatial attention during the delay period. For EEG Dataset 2 ([34], Experiment 1), each informative-cue trial began with a 100 ms cue (a thin bar extending from fixation) that pointed to the target’s location. After a 600 ms delay, a 200 ms search array appeared, consisting of a diamond target among seven square distractors. Participants responded by indicating whether the target was missing a left or right corner via a keypress. We only analysed informative-cue trials as uninformative cues presented a non-directional circle that lacked spatial specificity. EEG Dataset 3 ([34], Experiment 2) followed the same procedure as Dataset 2 but used a visually balanced cue to eliminate asymmetries. Instead of a single bar, eight bars radiated from fixation, with one bar in a different colour indicating the target location in the informative-cue condition. Again, we only analysed informative-cue trials because uninformative cues displayed eight bars of the same colour and did not induce a visuospatial attention effect. Across all three experiments, participants were instructed to maintain fixation throughout each trial and to blink shortly only after their response. Full details of the experimental procedures are reported in [33, 34].

For the TMS-EEG experiment [32], participants completed a selective attention task shown in Figure 3A. Participants completed two TMS-EEG sessions, each corresponding to one TMS condition (rh-TMS or ar-TMS) and consisting of 320 trials. In addition, each session included a no-TMS condition with 192 trials. Each trial began with a white fixation cross displayed for 500-1000 ms, followed by a 500 ms cue at the centre of the screen. The cue indicated both the attended location (‘<’ for left; ‘>’ for right) and the attended feature (‘c’ for colour; ‘o’ for orientation). TMS pulses were delivered during the first delay period for rh-TMS and ar-TMS conditions. The duration of the first delay varied based on the participant’s IAF in order to align the stimulus onset with the first or second alpha cycle peak. The first delay lasted (100 + 5000/IAF) ms or (100 + 6000/IAF) ms. After the first delay, the stimuli were presented on both sides of the screen for 100 ms, followed by a second delay of 1000 ms. Participants then saw a response screen and made their response within 5 seconds. In the colour task, they selected the target’s colour from a colour wheel, while in the orientation task, they adjusted a tilted bar to match the target’s orientation. Participants were instructed to fixate on the central cross throughout the experiment, except during the response period. More detailed procedures are available in Lu, et al. [32].

### TMS protocol

In the TMS-EEG experiment [32], participants first attended one MRI session to acquire a T1-weighted structural scan for neuronavagation. The stimulation site was the right intraparietal sulcus, with Montreal Neurological Institute (MNI) coordinates [x = 37, y = -56, z = 41], determined based on prior fMRI studies as the IPS part of the multiple-demand control network [41, 42]. We transformed these coordinates into each participant’s native space by normalizing their structural scan using the segment and normalize functions in SPM12. During the TMS-EEG sessions, we used the Brainsight 2 system for real-time neuronavigation, co-registering each participant’s head with their MRI scan to ensure precise targeting. We delivered TMS pulses using a DuoMag XT-100 stimulator with a figure-of-eight butterfly coil (DuoMag 70BF).

Before the first TMS-EEG session, we determined each participant’s resting motor threshold by identifying the lowest intensity that produced a visible muscle twitch in the contralateral abductor pollicis brevis in at least five out of ten pulses [43]. As shown in Figure 3A, participants received a train of five-pulse alpha rh-TMS or ar-TMS. Stimulation was at 110% of their motor threshold (63% of the maximal stimulator output on average), although the effective intensity was likely slightly lower due to attenuation of the magnetic field caused by the EEG cap. Stimulation began 100 ms after the first delay onset in each trial.

In the alpha rh-TMS condition, we applied pulses at each participant’s IAF. For the ar-TMS condition, we maintained the same total duration as rh-TMS but disrupted its rhythmic structure by fixing only the first and last pulse while randomly generating the remaining pulse timings.

In each session, we measured the pre-stimulus IAF in the first delay for each participant using the data of three EEG-only blocks (reported in [32]) before the TMS-EEG sessions. The pre-stimulus IAF was defined as the frequency with the maximum local power, calculated with fast Fourier transformation, in the alpha band (8-13 Hz) over a cluster of posterior electrodes around the right IPS (P2, P4, P6, P8, PO4, PO8, O2) during the first delay. In generating the arrhythmic trains, we ensured a minimum of 20 ms between pulses and monitored pulse intervals to prevent unintended entrainment. If more than three consecutive intervals fell within the alpha range (80–130 ms), we discarded and replaced the sequence with a new random pattern. Each TMS-EEG session (either rh-TMS or ar-TMS) included 1,600 pulses, with a minimum inter-train interval of 7.2 seconds. This protocol adhered to established TMS safety guidelines [44].

### EEG recording and preprocessing

We used preprocessed EEG data available from all four datasets (see details in [32–34]) and downsampled all recordings to 250 Hz for consistency. It should be noted that different EEG caps were used across datasets, leading to variations in electrode numbers and montage. EEG Dataset 1 [33] was recorded using 20 electrodes mounted in an elastic cap (Electro-Cap International, Eaton, OH). Electrodes were placed at International 10/20 sites: F3, FZ, F4, T3, C3, CZ, C4, T4, P3, PZ, P4, T5, T6, O1, and O2, along with five additional sites: OL, OR, PO3, PO4, and POz. EEG Dataset 2 and 3 [34] were recorded using 30 active electrodes mounted in an elastic cap (Brain Products). Electrodes followed the International 10/20 system, with signals recorded from Fp1, Fp2, F7, F3, Fz, F4, F8, FC5, FC1, FC2, FC6, C3, Cz, C4, CP5, CP1, CP2, CP6, P7, P3, Pz, P4, P8, PO7, PO3, PO4, PO8, O1, Oz, and O2. For analysis, we used 28 channels, excluding Fp1 and Fp2, as these sites were previously reported to have poor data quality due to excessive high-frequency noise and slow drifts [34].

For the TMS-EEG dataset [32], we recorded from 64 electrodes using TMS-compatible EEG equipment (actiCHamp Plus 64 System with actiCAP slim electrode cap, Brain Products UK). To aim TMS artefact removal, we used a sampling rate of 25,000 Hz during recording.

TMS-EEG preprocessing followed standardized pipelines implemented in EEGLAB [45], with additional processing using TMS-EEG Signal Analyzer (TESA) [46]. Preprocessing began by epoching the data from -1.1 to +2.1 s relative to the first TMS pulse and applying baseline correction using the fixation period (-1 to -0.6 s from the first TMS pulse). We removed the large TMS-induced artefact around each pulse (-2 to 10 ms) by setting these values to zero, and the data were then downsampled from 25,000 Hz to 1,000 Hz. We identified and removed bad trials and channels on an epoch-by-epoch basis using the TBT toolbox [47]. Next, we applied the first round of independent component analysis (ICA) to remove the large TMS-evoked muscle artefacts. After this, we extended artefact removal to -2 to +15 ms around each pulse and used cubic interpolation to reconstruct missing data. The data were then filtered using a band-pass filter (0.01–100 Hz) and a notch filter (48–52 Hz) (Butterworth method). In a second round of ICA, we removed remaining artefacts including eye movements, blinks, electrode artefacts, and residual muscle activity. Finally, we interpolated missing electrodes, re-referenced the data to the average, and re-epoched the data to stimulus onset. To match the other datasets, we then downsampled the data to 250 Hz. For the no-TMS condition, the sampling rate was 1,000 Hz during recording, and preprocessing steps were generally the same as in the TMS conditions but without TMS-specific steps (e.g., removal of TMS-induced artifacts and the first round of ICA).

### Irregular-resampling auto-spectral analysis (IRASA)

We conducted all analyses using MNE-Python [48], the Scikit-learn package [49], and custom Python scripts. Before applying IRASA, we removed time-locked event-related potentials (i.e., the average signal across trials) from each trial for each condition to isolate induced neural activity [50].

We used IRASA [35] to separate the oscillatory and the aperiodic components from the mixed power spectrum, using the delay period of each experiment as the analysis window (1000 ms for EEG Dataset 1, 600 ms for Datasets 2 and 3, and the 600 ms before stimulus onset for the TMS-EEG dataset). IRASA achieves this by resampling the EEG signals at pairwise non-integer rates (e.g., 1.1× to 1.9×, with corresponding inverse factors 0.9×–0.1×), which shifts the peak frequency of oscillatory components while leaving the broadband aperiodic component stable. To estimate the aperiodic component, we applied FFT-based power spectral analysis to each resampled signal and computed the geometric mean of the power spectra across resampling pairs, which minimises periodic components while preserving the broadband aperiodic activity. The median across all resampling factors was then taken to obtain the final aperiodic power spectrum. The oscillatory component was extracted by subtracting this aperiodic power spectrum from the original mixed power spectrum. Following standard recommendations [35], we used a set of resampling factors (*h*) from 1.1 to 1.9 in steps of 0.05 (as well as inverse factors from 0.9 to 0.1 in steps of 0.05). We estimated the power spectra in 1 Hz steps from 1 to 35 Hz, covering all relevant frequency bands with additional buffer areas, which help reduce edge effects and spectral leakage caused by the finite length of the analysis window [51]. For topographic plotting requiring baseline correction, we also applied IRASA during a baseline period before cue onset (600 ms for the three EEG datasets; 500 ms for the TMS-EEG dataset) using the same procedures as for the delay period.

To extract oscillatory alpha, aperiodic alpha, and mixed alpha signals, we computed power in the 8-13 Hz range from the oscillatory spectrum, aperiodic spectrum, and mixed power spectrum, respectively. For later correlation analyses, we also estimated aperiodic broadband power (3–30 Hz) by averaging power in this range from the aperiodic spectrum. Additionally, we computed aperiodic slope and intercept by fitting a linear function to the aperiodic power spectrum in log-log coordinates using least squares estimation.

### Multivariate decoding analysis

For each dataset, we performed multivariate decoding analysis on data from all EEG sensors after IRASA decomposition. To enhance the signal-to-noise ratio of decoding analysis, we grouped every four trials into “pseudo-trials” per condition by randomly selecting from the available trials [52]. We then standardized the data for each channel by subtracting the mean and dividing by the standard deviation across all epochs. We employed a 5-fold cross-validation procedure for classification using a linear support vector machine (SVM) for each signal of interest. In each iteration, pseudo-trials were randomly divided into five subsets, with four used for training and one for testing. This procedure was repeated five times, ensuring that all pseudo-trials contributed equally to both training and testing. The final decoding performance was obtained by averaging the results across these five iterations. We used balanced accuracy as the performance metric, defined as the arithmetic mean of sensitivity and specificity, as it provides a more robust evaluation for potentially imbalanced data [53]. To ensure that decoding results were not biased by a specific pseudo-trial selection, we repeated the entire procedure 25 times per participant and averaged the balanced accuracy over these repetitions for statistical testing.

### Lateralisation analysis

To qualitatively visualize lateralisation effects, we generated topographic plots for each dataset by subtracting EEG activity in the attend-right (3 o’clock) condition from the attend-left (9 o’clock) condition for each signal of interest. We applied baseline correction for each topographic plot by subtracting the baseline period data (before cue onset) after IRASA from the cue-period data after IRASA.

For quantitative analysis, we measured lateralisation effects using the modulation index (MI), a widely used metric for assessing lateralised neural activity [54–57]. The MI could be calculated separately for the left and right hemispheres, and we derived a combined MI by subtracting the right hemisphere MI from the left hemisphere MI to avoid possible bias in the analysis due to the hemisphere asymmetry. For each signal *S*, the MI was calculated as follows:

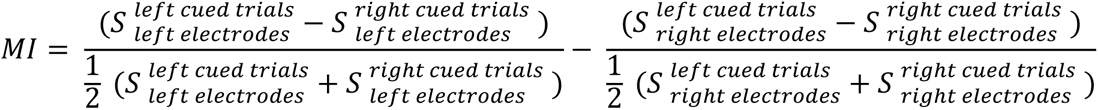

The amplitude of MI represents the strength of lateralisation, with larger absolute values indicating stronger lateralisation effects. The sign of MI reflects the direction of spatial modulation: Positive MI indicates that the signal decreases contralateral to the attended target, while negative MI indicates that the signal increases contralateral to the attended target. Given differences in electrode availability and montage across datasets, we selected posterior electrodes with slight variations. In EEG Dataset 1 [33], we used O1, P3, PO3, and T5 for the left hemisphere and O2, P4, PO4, and T6 for the right hemisphere. For the other three datasets, we used O1, P3, P7, PO3, and PO7 for the left hemisphere and O2, P4, P8, PO4, and PO8 for the right hemisphere.

### TMS modulation analysis

To assess TMS modulation effects, we generated topographic plots for each signal of interest using data after IRASA, computed by subtracting ar-TMS from rh-TMS. For quantitative analysis (Figure 3B), we measured modulation effects by averaging signals over a cluster of right posterior electrodes (P2, P4, P6, P8, PO4, PO8, O2, CP2, CP4, CP6, TP8, C2, C4, C6, T8) and computed the difference between rh-TMS and ar-TMS, following the channel selection from [32].

### Statistical analysis

All statistical analyses were performed in Python. For each dataset, we performed one-sample t-tests across participants to assess the significance of decoding performance, MI, and TMS modulation effects against their respective classification chance levels (0.125 for EEG datasets, 0.5 for the TMS-EEG dataset) or 0 (for MI and modulation effects). One-tailed t-tests were used for decoding analysis, while two-tailed t-tests were applied for MI and modulation analyses. To control the false discovery rate (FDR) across multiple comparisons, we applied the Benjamini-Hochberg (BH) procedure and reported results as significant at corrected *p* < 0.05. For comparisons across signals of interest, we used repeated-measures ANOVA for each dataset. In the TMS-EEG dataset, we also included TMS condition (rh-TMS vs. ar-TMS) as a factor. When ANOVA revealed a significant main effect, we performed post-hoc comparisons with Holm correction. For correlation analyses, we used Spearman’s correlation and applied FDR correction using the same BH procedure. For correlations between TMS-induced decoding changes and behavioural changes, we used one-tailed tests as we expected higher decoding to predict better behavioural performance. For all other correlations, we used two-tailed tests.

## Acknowledgements

This project was supported by UKRI MRC intramural funding (SUAG/093/G116768) to A.W., UKRI MRC Promote Units as National Assets award MC_PC_20046, and a Gates Cambridge Scholarship awarded to R.L. (OPP1144). For the purpose of open access, the author has applied a Creative Commons Attribution (CC BY) licence to any Author Accepted Manuscript version arising from this submission.

## Supplementary

**Figure S1.**
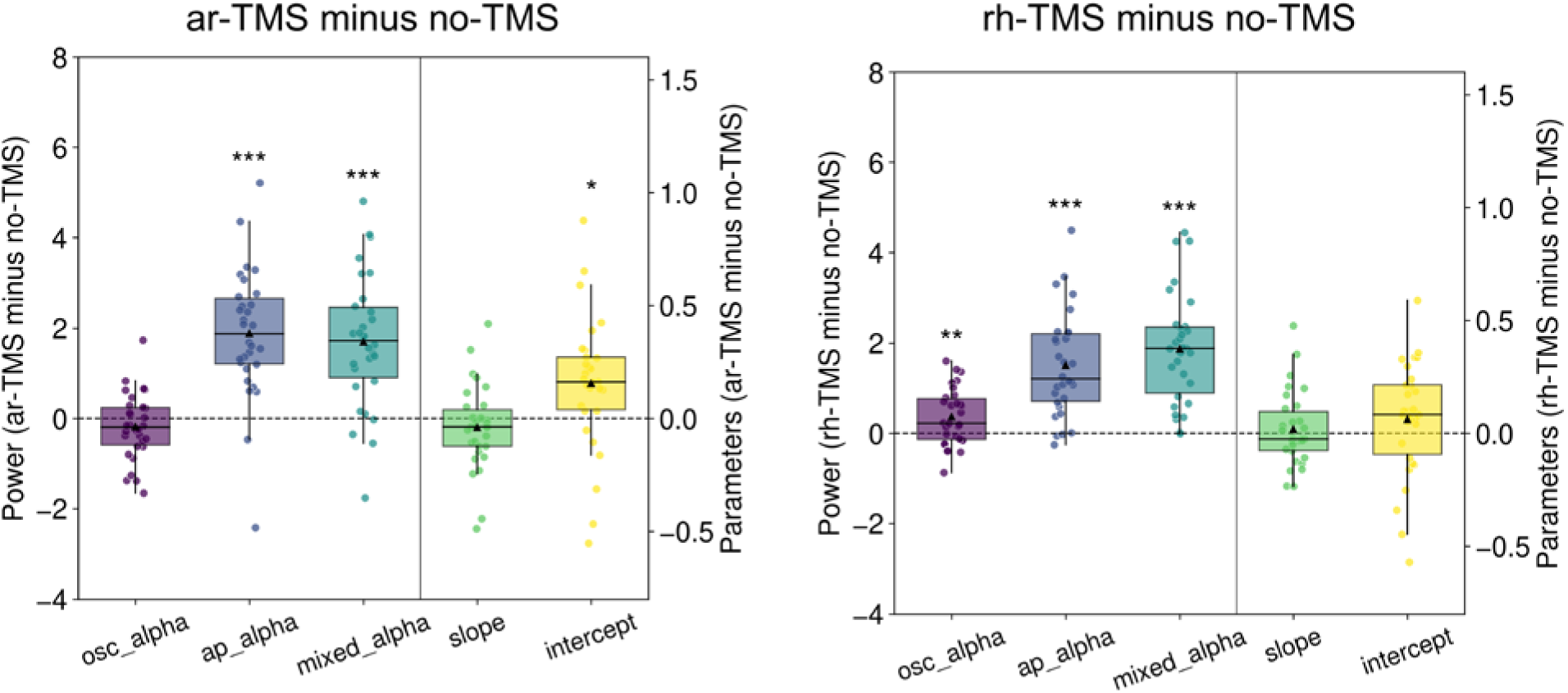
Comparison of rh-TMS and ar-TMS effects with no-TMS condition. TMS modulation effects for ar-TMS (left) and rh-TMS (right) relative to no-TMS condition during the delay period over right posterior regions. Box plots show activity differences (ar-TMS or rh-TMS minus no-TMS) for each type of signal. * *p* _FDR_ < 0.05, ** *p* _FDR_ < 0.01, *** *p* _FDR_ < 0.001

